# Development of two compatible plasmids to assess sRNA-mediated post-transcriptional regulation in *Acinetobacter baumannii*

**DOI:** 10.1101/2025.07.11.664372

**Authors:** Aalap Mogre, Orla Connell, Jessica White, Ali Shaibah, Karsten Hokamp, Fergal J. Hamrock, Kristina Schauer, Carsten Kröger

## Abstract

Post-transcriptional regulation can be mediated by small, regulatory RNAs in bacteria, which can act by base-pairing to a target messenger RNA. The discovery and mechanistic validation of base-pairing sRNAs in multidrug resistant *Acinetobacter baumannii* has been hampered by the lack of genetic tools to assess RNA-RNA interactions. Here, we created two compatible plasmids for *A. baumannii*, which addresses this need. The newly designed plasmids validated the known Aar sRNA-*carO* mRNA, and a new interaction of sRNA44 and the mRNA of the biofilm-associated protein Bap. The new plasmid system should accelerate the mechanistic characterisation of small, regulatory RNAs in *A. baumannii*.

**IMPACT STATEMENT:** Multi-drug resistance of pathogenic microorganisms is one of the greatest challenges for modern medicine. Carbapenem-resistant *Acinetobacter baumannii* are considered a highly critical organism, yet we are only beginning to understand its physiology and mechanisms of gene regulation. Post-transcriptional regulation by base-pairing, small RNAs is an understudied area, partly because of the lack of genetic tools to investigate them. In this study, we developed a 2-plasmid system to assess sRNA-mRNA interactions, which will greatly accelerate the discovery and validation of small, regulatory RNAs and their target molecules.

**DATA SUMMARY:** Plasmid sequences of pAMCK14-sRNA44 and pAMCK18-Bap have been made available in GenBank of National Center for Biotechnology Information (accession numbers PV916437 and PV916438).

## INTRODUCTION

Small regulatory RNAs (sRNAs) are ubiquitous post-transcriptional regulators in bacteria [1, 2]. Many of them function by base-pairing to another RNA molecule, typically a messenger RNA (mRNA) molecule, to modulate its translation [3]. One of the challenges in determining the biological function of base-pairing regulatory sRNAs is identifying their target mRNA. Methods to identify RNA-RNA interactions range from bioinformatic predictions to several experimental approaches that ideally should be combined [4]. Several methods have been developed to experimentally verify RNA-RNA interactions, including high-throughput, proximity ligation-based methods such as RIL-seq, Hi-GRIL-seq, iRIL-seq or CLASH [5, 6]. As these methods rely on proximity ligation, which can ligate transiently interacting RNAs that do not represent biologically relevant RNA-RNA pairs, RNA-RNA interactions identified by high-throughput methods must be independently validated to distinguish biologically relevant from transient, less relevant interactions.

A genetic system consisting of two replication-compatible plasmids has been successfully used to validate and dissect mechanistic details of sRNA-mRNA interactions in *Enterobacteriaceae* [7–9]. The sRNA is expressed from one plasmid and is assessed whether it alters the translation of the target mRNA-green fluorescent protein (GFP or superfolder GFP) reporter gene fusion expressed from the second plasmid [7, 10, 11]. Introduction of nucleotide changes to disrupt and restore interacting sRNA-mRNA duplexes allows precise identification of nucleotides essential for base-pairing. The original 2-plasmid system has been instrumental to characterise many regulatory sRNAs in *E. coli* and *S. enterica*. However, these original plasmids are not always compatible with (co-)replication and selection in other Gram-negative bacteria, hindering the progress of studying sRNA mediated post-transcriptional regulation in non-model or multidrug resistant organisms. For example, these plasmids are not replicative in *A. baumannii* where much less is known about the regulatory functions and biological roles of sRNAs [12].

Recently, it was demonstrated that plasmids harbouring the pWH1266 and pRSF1010 origins of replication can be stably co-maintained in *Acinetobacter* [13, 14]. Arguably, the most used plasmid in *A. baumannii* is pWH1266, which is a shuttle plasmid constructed from fusion of a natural *Acinetobacter calcoaceticus* plasmid pWH1277 and pBR322 [15]. The pWH1266 origin of replication was subsequently widely used to generate plasmids for gene overexpression or chromosomal integration in *A. baumannii* under the control of various regulatory systems [14, 16–19]. Plasmids with a pRSF1010 origin of replication support a broad host range and were shown to replicate in *Acinetobacter* spp. decades ago [20]. To create a 2-plasmid system for assessing sRNA-mRNA interactions in *A. baumannii*, we aimed to generate two plasmids carrying the replication compatible pWH1266- and pRSF1010-derived origin of replications. The plasmids were evaluated for their ability to replicate and functionally substitute for the historical pP_L_- and pXG-plasmids created for *Enterobacteriaceae* using the same constitutive P_LtetO_ and P_LlacO_ promoters and a truncated sfGFP to create translational fusions in *A. baumannii* [7, 10, 11].

## METHODS

### Bacterial strains, growth conditions and transformations

*Escherichia coli* TOP10 and *Acinetobacter baumannii* AB5075 were maintained on lysogeny (L-) broth agar plates (Lennox, 10 g/l tryptone, 5 g/l yeast extract, 5 g/l NaCl, 15 g/l agar) at 37°C. Antibiotics (tetracycline (12 μg/ml), apramycin (60 μg/ml)) were added when required. *E. coli* cells were transformed by electroporation or heat-shock transformation. *A. baumannii* AB5075 cells were transformed by electroporation.

### Chemical transformation

*E. coli* TOP10 cells were made chemically competent using the TSS method [21]. A 4-ml overnight culture was set up from a single colony and incubated at 37°C, 220 rpm. The next day the culture was diluted 1:100 in 50 ml L-broth in a 250 ml Erlenmeyer flask. The flask was incubated at 37°C, 220 rpm until A_600_ reached 0.6. Cells were pelleted at 4000 rcf, 4°C, 15 minutes. Supernatant was discarded and cells were resuspended in 2 ml TSS solution (L-broth containing 10% (w/v) polyethylene glycol 3350, 5% (v/v) Dimethyl sulfoxide (DMSO), 50 mM MgCl_2_ at pH 6.5). TEDA reactions were incubated with 100 µl TSS competent TOP10 cells on ice for 30 min. Heat shock was performed at 42°C for 45 s in a water bath. Cells were incubated on ice for 2 min. 800 µl L-broth was added and cells were incubated on a Thermo mixer at 37°C, 600 rpm, for 1 h when selecting on apramycin and 1.5 h when selecting on tetracycline. Following recovery, cells were pelleted at 10,000 rcf, resuspended in 50 µl L-broth and plated on agar plates containing antibiotic(s).

### Electroporation

*E. coli* (electroporation used only for the three fragment TEDA cloning reaction): To make electrocompetent *E. coli* TOP10 cells, a 4-ml overnight culture was set up from a single colony and incubated at 37°C, 220 rpm. The next day the culture was diluted 1:100 in 50 ml L-broth in a 250 ml Erlenmeyer flask. The flask was incubated at 37°C, 220 rpm until A_600_ reached 0.6. Cells were pelleted at 4000 rcf, 4°C, 15 min. Supernatant was discarded, and cells were then washed once with 1 ml sterile ice-cold water and once with 1 ml sterile ice-cold 10% (v/v) glycerol with pelleting steps being carried out at 10,000 rcf, 4°C, 2 min. Cells were finally resuspended in 500 µl sterile ice-cold 10% glycerol thus concentrating the cells 100-fold. The desalted TEDA reaction was mixed with 50 µl competent cells in a 2-mm electroporation cuvette. Electroporation was performed at 2.5 kV with 200 Ω resistance and 25 µF capacitance. Recovery was carried out as above. To make electrocompetent *A. baumannii* AB5075 cells, a 4-ml overnight culture was set up from a single colony and grown for 16 h at 37 °C, 220 rpm. Cells were pelleted at 4000 rcf, 4°C, 15 min. Supernatant was discarded, and cells were then washed once with 1 ml sterile ice-cold water and once with 1 ml sterile ice-cold 10% glycerol with pelleting steps being carried out at 10,000 rcf, 4°C, 2 min. Cells were finally resuspended in 400 µl sterile ice cold 10% glycerol thus concentrating the cells 10-fold. 25-50 ng plasmid DNA was mixed with 50 µl competent cells in a 2-mm electroporation cuvette. Electroporation and recovery were carried out as above.

### Plasmid constructions

Oligonucleotides used for priming PCRs are listed in **Table SX**. Plasmids were assembled from PCR products using Seamless Ligation Cloning Extract (SLiCE) or T5 exonuclease DNA assembly (TEDA) [22, 23]. Plasmid inserts were confirmed by Sanger sequencing (Eurofins) and whole plasmids pAMCK14-sRNA44 and pAMCK18-Bap were sequenced by Azenta (Leipzig, Germany).

### Measurement of fluorescence

A single colony was inoculated in 1 ml L-broth containing apramycin and tetracycline and grown for 30 h at 37°C and 220 rpm. Apart from the low volume of broth used to set up the cultures, tubes were also placed in the incubator shaker at an angle to improve aeration. This was necessary for a better production of fluorescence signal from the expressed sfGFP in *Acinetobacter baumannii* AB5075. 100 μl of the culture was then transferred into the wells of a white 96-well plate with clear bottom (Corning, Costar) and fluorescence (arbitrary units) and absorbance at 600 nm (A_600_) were measured in a BioTek Synergy HTX Multimode Reader. Fluorescence was measured with a top read, with excitation filter 485/20, emission filter 528/20, and gain 45. The fluorescence of different strains was compared by quantifying the fluorescence observed relative to A_600_ (F/A_600_). L-agar plates were imaged using epi blue light and a 510DF10 filter in the ImageQuant LAS4000 (GE Healthcare).

## RESULTS

### Construction of plasmids to study sRNA-mRNA interactions

The original 2-plasmid system comprises the pP_L_ plasmid which carries the sRNA gene and the pXG10/pXG10sf plasmid which carries the target mRNA-*GFP* fusion [7, 11]. These plasmids contain origin of replications (ColE1 and pSC101*) and selection genes (resistance to ampicillin and chloramphenicol) that are unsuitable for use in multi-drug resistant *A. baumannii* strains, such as the widely used strain AB5075 [24]. To adapt the system for *A. baumannii* (**Figure 1**), the sRNA and mRNA expression regions of the original 2-plasmid system were retained and origins of replication and resistance cassettes suitable for *A. baumannii* were introduced. To construct the sRNA expression plasmid (pAMCK14, **Figure 1**), a fragment of the pP_L_ plasmid containing the P_LlacO_ driven sRNA-expression region was fused with the *rrnBT1* terminator and the ColE1 origin of replication and combined with a fragment of the pAMCK2 plasmid carrying the widely used pWH1266 origin of replication and the apramycin resistance gene, *aacC4*, from pMHL2 [15, 25]. The ColE1 origin of replication enables replication in *E. coli*, while the pWH1266 origin of replication allows replication in *A. baumannii*, making the resulting plasmid a shuttle vector. To construct the control plasmid (pAMCK15, **Figure 1**), the sRNA gene was deleted from pAMCK14 while retaining the *rrnB1* terminator. This plasmid is equivalent to the pJV300 control plasmid used in the original 2-plasmid system and would express a short nonsense RNA [7]. To construct the target mRNA expression plasmid, a fragment from pXG10sf containing the P_LtetO_-driven target mRNA region fused to the start codon-lacking *sfGFP*, an *rrnBT1* terminator and the pSC101 origin of replication was combined with a fragment containing the pRSF1010 origin of replication which was obtained from *S. enterica* serovar Typhimurium SL1344 [26], and a fragment of the pBR322 plasmid carrying the tetracycline resistance gene *tetA*. The resulting 10.3 kb plasmid contains both the pSC101 and pRSF1010 origins of replication and presents a shuttle vector capable of replication in both *E. coli* and *A. baumannii* AB5075. Although fluorescent, mutations in *sfGFP* were detected upon sequencing, which were corrected. Because the pRSF101 origin of replication works in *A. baumannii* and *E. coli*, the pSC101 origin of replication was removed to construct the final plasmid (pAMCK18, **Figure 1**) which retains only the P_LtetO_-driven target mRNA-expression region fused to *sfGFP* and terminated by *rrnBT1*. This ∼8 kb pAMCK18 plasmid is also a shuttle vector capable of replication in both *E. coli* and *A. baumannii* as expected from the broad range pRSF1010 origin of replication. The retention of the sRNA and target mRNA expression regions of the original 2-plasmid system ensured that the primers used to amplify the backbones and inserts for cloning could be used with the original and new 2-plasmid systems.

**Figure 1.**
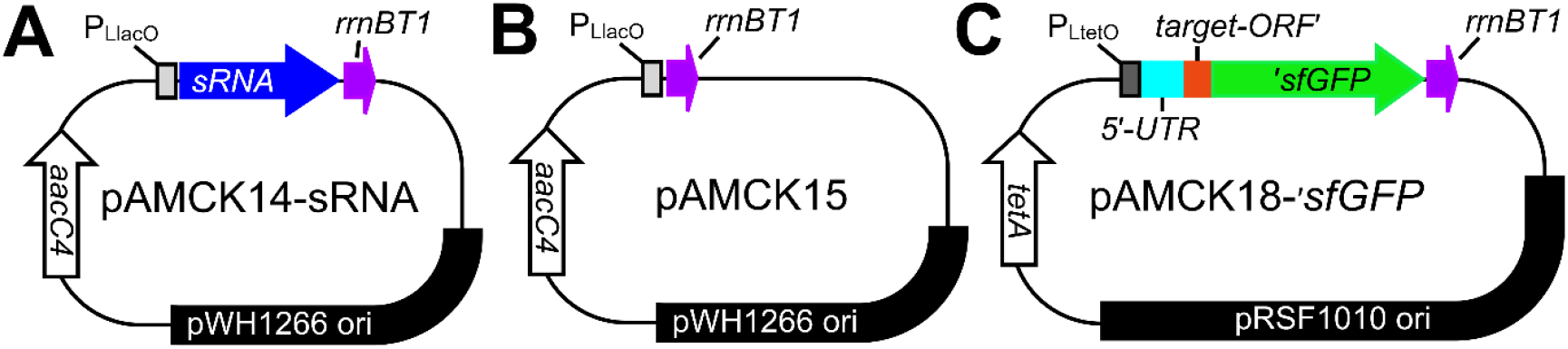
Schematic plasmid maps of pAMCK14 **(A)**, pAMCK15 **(B)** and pAMCK18 **(C)**. The plasmid pAMCK14 **(A)** expresses a small RNA from the P_LlacO_ promoter, which is terminated by the rrnBT1 terminator. In pAMCK15 **(B)**, the same P_LlacO_ promoter is terminated almost immediately and expresses a small, non-sense RNA, which is identical with the non-sense RNA expressed from pJV300 [7]. Both plasmids possess the pWH1266 origin of replication and the *aacC4* gene from pMHL-2 conferring resistance to apramycin [25]. Plasmid pAMCK18 **(C)** expresses a translational fusion of a target gene fused to a start-codon-less sfGFP obtained from pXG10sf [11]. The fusion protein is expressed from the P_LtetO_ promoter and terminated by the rrnBT1 terminator as in the original pXG10 plasmid [7, 11]. Plasmid pAMCK18 harbours the *tetA* gene conferring tetracycline resistance and the pRSF101 origin or replication. The plasmid schematics are not drawn to scale.

### Validation of the Aar-*carO* translational repression in *E. coli* TOP10 and *A. baumannii* AB5075

To evaluate the functionality of the new 2-plasmid system in *A. baumannii* AB5075 and *E. coli* TOP10, translational repression of the *carO* mRNA by the sRNA Aar of *A. baumannii* AB5075 was assessed. The interaction was previously validated in *E. coli* using the original 2-plasmid system where expression of Aar from the pP_L_ plasmid reduced fluorescence from the pXG10-carO-sfGFP reporter compared to the pJV300 control, however, validation of the interaction in *A. baumannii* AB5075 previously required a more complex strategy relying upon the introduction of several chromosomal mutations [27]. Identical sequences of Aar and *carO* used in original 2-plasmid system were cloned into pAMCK14 (resulting in pAMCK14-Aar expressing Aar) and pAMCK18 (resulting in pAMCK18-carO expressing the CarO-sfGFP fusion protein). As a control, pAMCK18-carO was paired with the control plasmid pAMCK15. Plasmid combinations were measured for fluorescence in *E. coli* TOP10 and *A. baumannii* AB5075 (Δ*aar* when containing both plasmids) and normalised to culture absorbance (**Figure 2**). In *E. coli*, the new system largely recapitulated the findings from the original 2-plasmid system (**Figure 2A**). In *E. coli*, the level of fluorescence of the pAMCK18-carO fusion was ∼1.5-fold higher than the pXG10sf-carO variant possibly due to the corrected mutations in *sfGFP*. Co-expression of wild-type Aar reduced translation of *carO* mRNA by ∼77%, comparable to ∼90% reduction observed with the original system (pXG10sf/pP_L_/pJV300) showing that the new 2-plasmid system is functionally equivalent to the original system. The level of fluorescence in *A. baumannii* AB5075 Δ*aar* pAMCK18-carO expressing wild-type Aar from pAMCK14 was reduced (p = 0.0002) compared the strain harbouring the control plasmid pAMCK15, validating translational repression of CarO-sfGFP in *A. baumannii* AB5075 ((**Figure 2B**), [27]). Consistent with previous observations [27], the repression of *carO* translation by Aar is much weaker in *A. baumannii* compared to *E. coli*, suggesting that there are additional endogenous regulatory factors influencing the level of *carO* translation using the 2-plasmid system in *A. baumannii*. Taken together, the data suggest that the new 2-plasmid system enables the study of sRNA-mediated post-transcriptional regulation in *E. coli* and *A. baumannii*.

**Figure 2.**
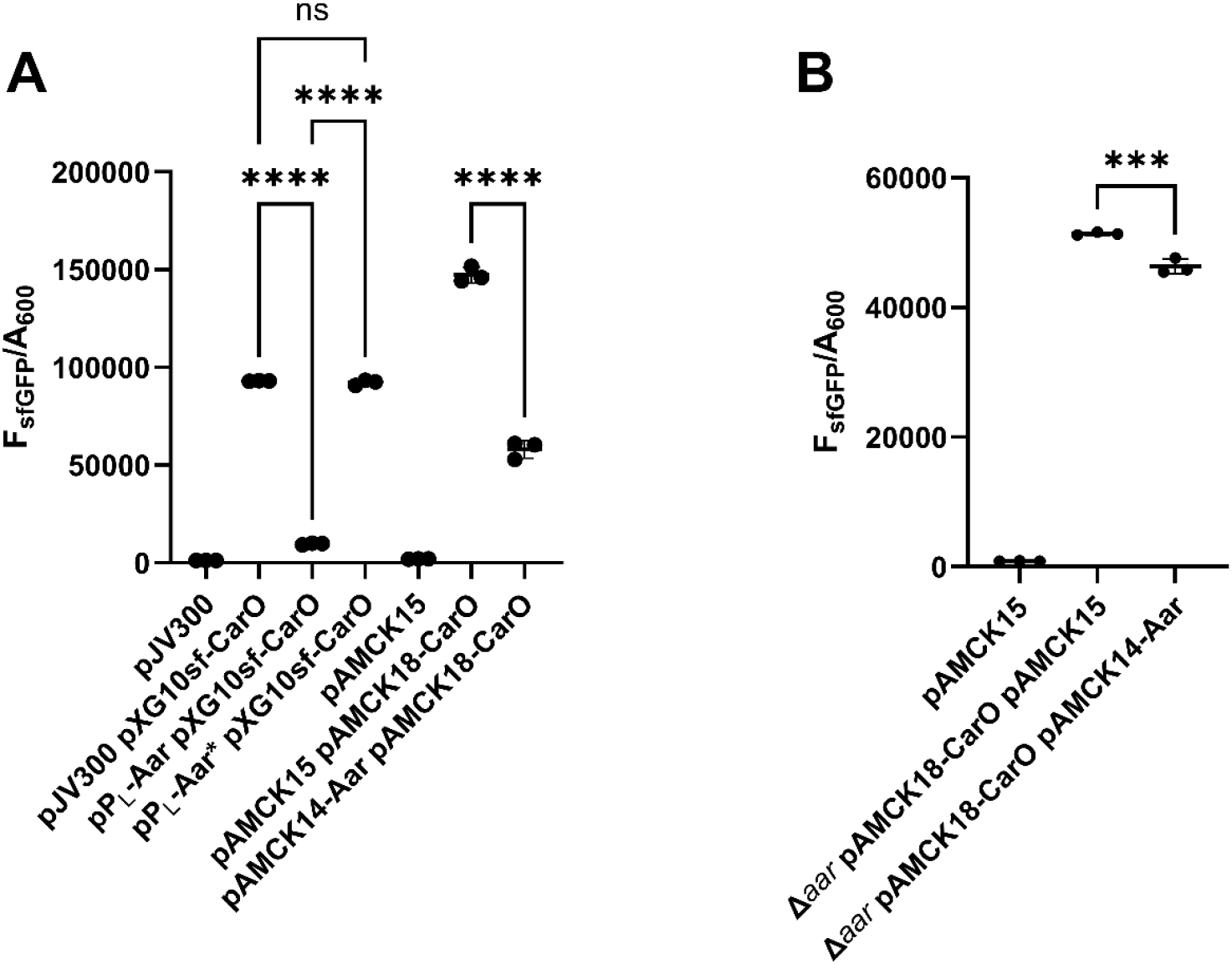
The small RNA Aar represses translation of CarO′-sfGFP in *E. coli* TOP10 **(A)** and *A. baumannii* AB5075 **(B)**. Comparison of original 2-plasmid system consisting of pJV300, pPL and pXG10sf plasmids and newly developed pAMCK14, pAMCK15 and pAMCK18 system showing that expression of Aar represses translation of CarO′-sfGFP validating earlier findings [27]. The data are expressed as fluorescence (F_sfGFP_) divided by culture absorbance (A_600_) after 30 h. Three independent experiments were performed per strain. Whether differences are statistically significant was assessed by One-Way ANOVA with Tukey’s multiple comparisons test. *** indicates *P*_adj_ < 0.001, **** indicates *P*_adj_ < 0.0001, ns indicates *P*_adj_ > 0.05.

### Translation of *bap* is inhibited by sRNA44

To test a novel, unvalidated sRNA-mRNA interaction of *A. baumannii*, a putative sRNA-mRNA pair was selected which had been previously identified by Hi-GRIL-seq [27]. Biofilm formation is a key virulence trait of *A. baumannii* virulence and persistence in hospital environments, and the biofilm associated protein Bap is involved in biofilm formation and is widely present in *Acinetobacter* spp. [28, 29]. Hi-GRIL-seq recovered 28 chimeric sequencing reads ligated between sRNA44 and *bap* mRNA, suggesting that sRNA44 may regulate the translation of *bap* mRNA through base-pairing [27]. Two additional mRNAs were ligated to sRNA44 encoding a hypothetical gene (*ABUW_RS05340*, 22 chimeric reads) and the type-I-F CRISPR-associated helicase Cas3 (11 chimeric reads, **Figure 3A**, [27]). The *bap* mRNA was prioritised because sRNA44-*bap* mRNA chimeric reads were most frequent and IntaRNA predicted the formation of an RNA duplex between sRNA44 and *bap* mRNA just downstream of the start codon of the *bap* mRNA (**Figure 3B**, [30]). Chimeric reads from all Hi-GRIL-seq conditions mapped to the AB5075 genome (IND = T4 RNA ligase induced, IMIP = T4 RNA ligase induced with additional brief “shock” with sub-inhibitory concentration of imipenem, DIP = T4 RNA ligase induced with additional brief limitation of iron by the addition of 2,2′-dipyridyl, NI = control, T4 RNA ligase not induced) matched the predicted duplex location (**Figure 3C**). The Hi-GRIL-seq data for sRNA44 can be accessed and browsed online using Jbrowse2: https://bioinf.gen.tcd.ie/hi-gril-seq/sRNA44 [31]. To test the interaction, plasmids pAMCK14-sRNA44 (expressing sRNA44) and pAMCK18-Bap (expressing the *bap* 5′ UTR fused to sfGFP, including the predicted duplex region and the first thirteen codons of *bap*) were constructed (**Figure 3C**). The transcriptional start sites for sRNA44 and *bap* were previously defined by differential RNA-seq [32]. For comparison, the original 2-plasmid system was constructed with sRNA44 expressed from pP_L_-sRNA44 and the Bap-sfGFP fusion protein expressed from pXG10sf-Bap. To disrupt the interaction of sRNA44 with *bap* mRNA, nucleotides 35-38 of sRNA44 were mutated from 5′-CAGG-3′ to 5′-GACC-3′ in pPL-sRNA44 to construct pPL-sRNA44* and in pAMCK14 to construct pAMCK14-sRNA44* (**Figure 3B**). RNA structure predictions of sRNA44 and sRNA44* showed that the mutated nucleotides are located in a single-stranded region and did not interfere with their structure (**Supplementary Figure 1**). Plasmid combinations were again measured for fluorescence as before and normalised on culture absorbance. Using the original system in *E. coli*, expression of sRNA44 from pP_L_-sRNA44 led to a reduction of fluorescence by 77% compared to the strain carrying the pJV300 control plasmid indicating that sRNA44 represses production of the Bap-sfGFP fusion protein (**Figure 4A**). Introduction of the three point mutations of sRNA44* relieved the repression and Bap-sfGFP abundance was even slightly higher than the strain containing the pJV300 control plasmid. In both *E. coli* TOP10 (78% reduction of fluorescence, **Figure 4A**) and *A. baumannii* AB5075 (51% reduction of fluorescence, **Figure 4B** and **4C**), co-expressing pAMCK14-sRNA44 along with pAMCK18-Bap resulted in a decrease in fluorescence signal compared to the control plasmid pAMCK15.

**Figure 3.**
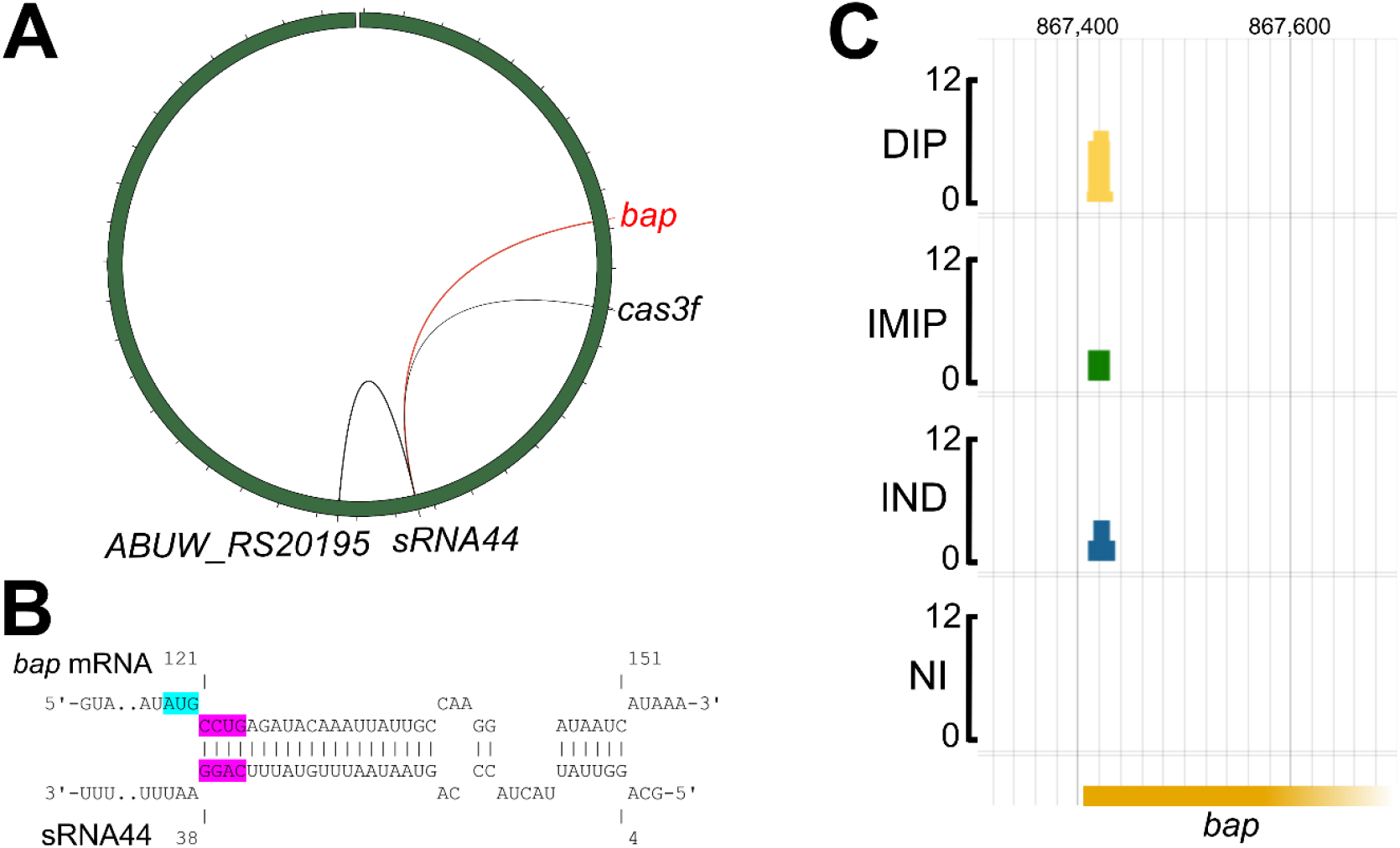
**(A)** Three mRNAs (*bap, ABUW_RS20195* and *cas3f*) were ligated to sRNA44 in a Hi-GRIL-seq experiment [27]. The green circle represents the chromosome of *A. baumannii* AB5075 with the origin of replication at the top of the circle and the location of *sRNA44, bap, ABUW_RS20195* and *cas3f* are highlighted. The width of the lines is proportional to the number of chimeric sequencing reads obtained from the Hi-GRIL-seq experiment. **(B)** Predicted interactions of *bap* mRNA (top sequence) with sRNA44 (bottom sequence) using IntaRNA [30, 54]. The *bap* start codon is highlighted in light blue. The nucleotides 35-38 in sRNA44 were mutated from 5′-CAGG-3′ to 5′-GACC-3′ to create sRNA44* (highlighted in pink with their base-pairing nucleotides in *bap* mRNA). The numbers indicate positions relative to the transcriptional start sites. **(C)** Mapped Hi-GRIL-seq derived chimeric reads consisting of bap mRNA and sRNA44. The scale is 0-12 reads. The numbers at the top of the figure indicate the genomic location. The *bap* gene is truncated for illustration purposes. IND = T4 RNA ligase induced, IMIP = T4 RNA ligase induced with additional brief “shock” with sub-inhibitory concentration of imipenem, DIP = T4 RNA ligase induced with additional brief limitation of iron by the addition of 2,2′-dipyridyl, NI = control, T4 RNA ligase not induced

**Figure 4.**
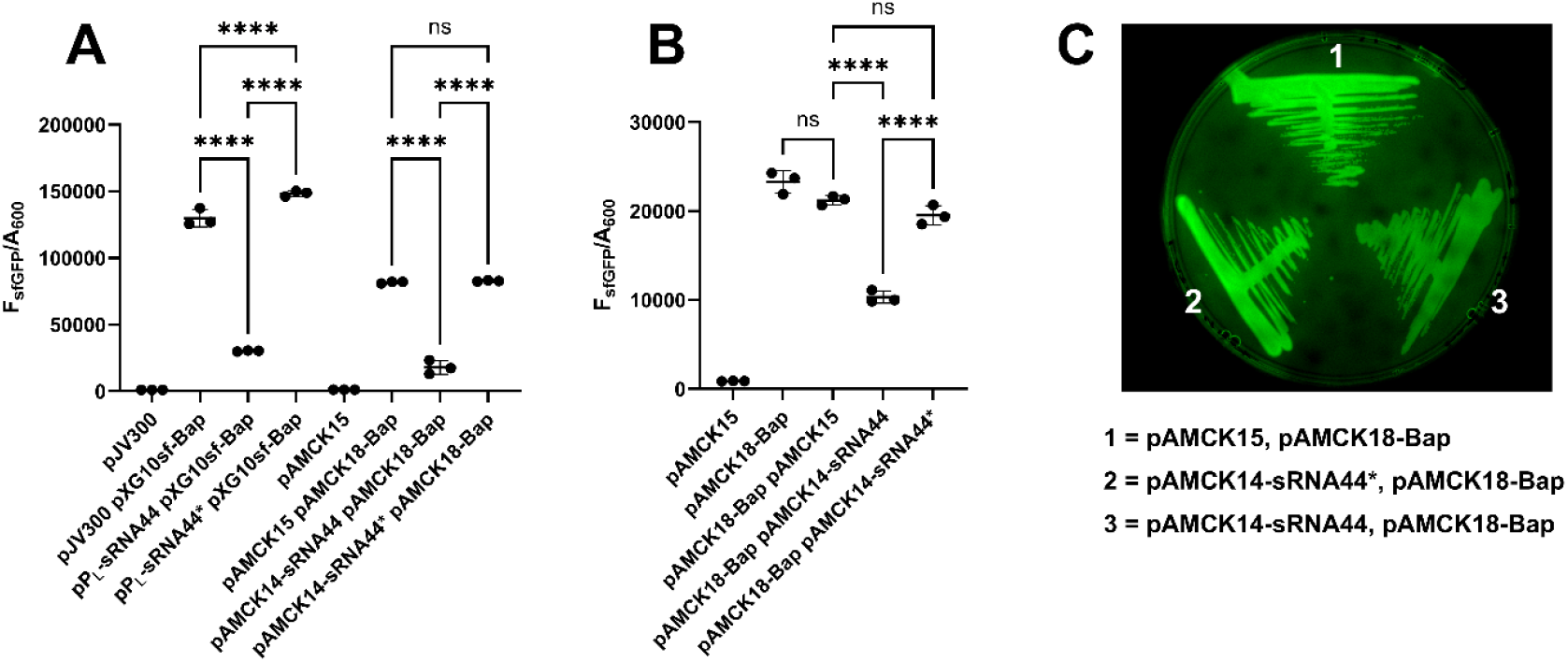
The small RNA sRNA44 represses translation of Bap′-sfGFP in *E. coli* TOP10 **(A)** and *A. baumannii* AB5075 WT (B and C). The data are expressed as fluorescence (F_sfGFP_) divided by culture absorbance (A_600_) after 30 h **(A and B)**. Three independent experiments were performed per strain. Whether differences are statistically significant was assessed by One-Way ANOVA with Tukey’s multiple comparisons test. **** indicates *P*_adj_ < 0.0001, ns indicates *P*_adj_ > 0.05. In (C), *A. baumannii* AB5075 carrying the indicated plasmids was streaked on an L-agar plate containing apramycin and tetracycline and grown for 16 h before the plate was imaged.

In contrast to the sfGFP fusions with CarO, the Bap-sfGFP abundance was higher in the pXG10sf-Bap compared to pAMCK18-Bap in *E. coli* (**Figure 2A, 4A**). In both bacteria, co-expressing the mutant variant sRNA44* from pP_L_-sRNA44* or pAMCK14-sRNA44* along with pAMCK18-Bap increased fluorescence was observed compared to the strains expressing wild-type sRNA44 suggesting that the nucleotides 35-38 of sRNA44 are required for base-pairing (**Figure 4B**).

## DISCUSSION

Small, regulatory RNAs which base-pair with a target RNA molecule are found in all kingdoms of life and provide an additional layer of gene regulation. In bacteria, foundational mechanistic understanding of these interactions was first generated in *E. coli* and *S. enterica* by using genetic tools designed for these model organisms. One versatile and highly utilised genetic tool to study the mechanisms of sRNA-mediated post-transcriptional regulation consists of two plasmids of which one expresses a small RNA and the other the target RNA fused to a reporter gene, first GFP and later sfGFP [7, 7, 11, 33–43]. This general approach has since been adopted to other bacteria, such as *Vibrio cholerae* and *Klebsiella pneumoniae* [44–47] and used to probe sRNA-mRNA interactions in *Bacteroides thetaiotaomicron* and *A. baumannii* using *E. coli* as a heterologous host [27, 48]. The key advantages of this system are that the sRNA and mRNA are expressed from separate plasmids, enabling clean and rapid validation of *trans* interactions and that plasmids are easily constructed and quickly mixed and matched. In this study, the original 2-plasmid system was extended to function in multidrug resistant *A. baumannii* by swapping origins of replications with ones that support stable replication in *A. baumannii* and suitable selection genes. Both plasmids still contain origins of replication that function in *E. coli* to allow the study of RNA-RNA interactions in a heterologous host. We note that fluorescence in *E. coli* is ∼3-4-fold higher from the same pAMCK18-carO/bap plasmids compared to *A. baumannii*, which could be due to a lower transcription or plasmid copy number in *A. baumannii*. Alternatively, host-specific differences in the availability or function of RNA-binding proteins, such as the notable differences of Hfq between *E. coli* and *A. baumannii* [49– 51], may modulate the stability or formation of sRNA–mRNA duplexes in the native versus heterologous background. Nevertheless, we validated the translational repression of CarO by the sRNA Aar in *A. baumannii* [27], and validated a new sRNA-mediated regulation of an *A. baumannii* virulence factor, the Bap protein [28, 29]. Future experiments will be required if the translational repression of Bap by sRNA44 has physiological consequences such as the level of biofilm formation or cellular adhesion. Beyond validating native regulatory interactions, the newly constructed 2-plasmid system may also be used to construct and benchmark synthetic small RNAs in *A. baumannii*, similar to toolboxes created for *E. coli* [52, 53].

Although the utility of the pAMCK 2-plasmid system in *A. baumannii* is clear, a few limitations and challenges should be acknowledged. Because the use of two multi-copy plasmids and constitutive promoters, the stoichiometry of sRNA-mRNA interactions and associated effects on cellular physiology will be different from wild-type cells which necessitates follow-up experiments to validate biological relevance. Differences in fluorescence produced by protein fusions are not easily anticipated due to inherent individual folding kinetics of every protein under assessment and host-specific variation in translation initiation of target protein-GFP fusions may also limit the detectability of fluorescence. In cases where the signals fall below the detection limit, sfGFP abundance can still be measured by Western blotting using GFP-specific antibodies, though this is more laborious. Because sRNAs and target genes must be cloned at nucleotide resolution, SLIC, SLiCE, TEDA, Gibson assembly or similar methods of cloning are required. Because this involves PCR-based amplification of the plasmid backbones, we recommend whole plasmid sequencing after cloning which multiple sequencing providers offer at low cost. Finally, although P_LlacO_ and P_LtetO_ are theoretically inducible in strains containing *lacI* and *tetR*, such regulatory elements are not yet standard in *A. baumannii*. Incorporation of *lacI* and *tetR* into the host would broaden the utility of the system when investigating regulatory effects following induction of sRNAs [10].

In conclusion, the newly developed shuttle pAMCK 2-plasmid system enables robust validation of sRNA-mRNA interactions facilitates mechanistic dissection of the interactions through targeted mutagenesis of predicted base-pairing regions in both *A. baumannii* and *E. coli*.

## ACKNOWLEDGEMENTS

We thank Xavier Charpentier (Inserm/CIRI, Lyon, France) and Jörg Vogel (Helmholtz Institute for RNA-based Infection Research, Würzburg, Germany) for sharing plasmids. Deirdre Muldowney (TCD) is acknowledged for technical support.

## AUTHOR CONTRIBUTIONS

A.M. and C.K. designed the study. A.M., O.C., J. W., A. B., F.J.H., K.S. and C.K. performed experiments. A.M., F.J.H., K.S. and C.K. analysed the data and prepared figures. A. M. and C.K. wrote the original draft. O.C., J. W., A. B., F.J.H., K.S. reviewed and edited the draft.

