## Supplementary data for "Development of two compatible plasmids to assess sRNA-mediated post-transcriptional regulation in *Acinetobacter baumannii*"

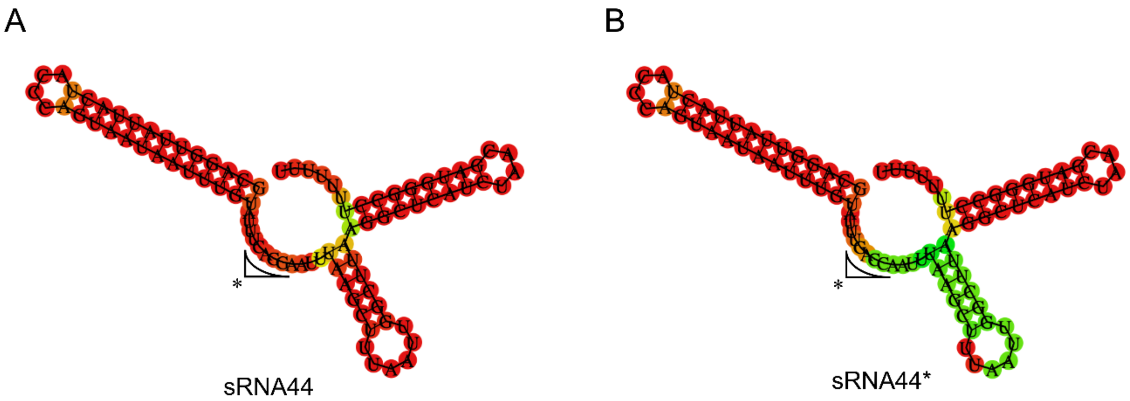


**Figure S1.** Structure predictions of sRNA44 **(A)** and sRNA44* **(B)**. The prediction was carried out using RNAfold [55]. The positions of the mutated nucleotides are highlighted with an asterisk.

**Table S1.** List of DNA oligonucleotides used in this study.

| No. | Purpose | Sequence (5′-3′) |
|---|---|---|
| 1438 | Amplification of sRNA44 insert | GTGAGCGGATAACAAGATACTGAGCACGCAGGTTATTACTACCCAG |
| 1439 | Amplification of sRNA44 insert | GCCTTTCGTTTTATTTGATGCCTCTAGACACCTATTATTATCAAGTGGC |
| 143 | Amplification pXG10sf backbone | GCTAGCGGATCCGCTGGCTCCGCTGC |
| 134 | Amplification pXG10sf backbone | ATGCATGTGCTCAGTATCTCTATCAC |
| 141 | Amplification $pP_L$ backbone | GTGCTCAGTATCTTGTTATCCGCTCAC |
| 142 | Amplification $pP_L$ backbone | TCTAGAGGCATCAAATAAAACGAAAGGC |
| 1436 | Amplification of bap insert | ACTGAGCACATGCATGTAAAAATGATCACATTAC |
| 1437 | Amplification of bap insert | AGCGGATCCGCTAGCTTTATGATTATCCTTGGC |
| 2517 | Construction of pAMCK14 (pPL-sRNA44-AbOri-aacC4) | GGACTTATCATCCAACCTGTCTGTCAGACCAAGTTTACG |
| 1958 | Construction of pAMCK14 (pPL-sRNA44-AbOri-aacC4) | TTTCCCCGAAAAGTGCCACCTG |
| 2518 | Construction of pAMCK14 (pPL-sRNA44-AbOri-aacC4) | CGTAAACTTGGTCTGACAGACAGGTTGGATGATAAGTCC |
| 2519 | Construction of pAMCK14 (pPL-sRNA44-AbOri-aacC4) | TAATGGTTTCTTAGACGTCAGGTGGCACTTTTCGGGGAAAGATCGTAGAAATATCTATG |
| 2520 | Construction of pAMCK12 (pXG10-bap-pRSF1010-ori-tetA) | AGAACATATCCATCGCGTCCGCCATCTCCAGCAGCCACGTCTCATTTTCGCCAGATATC |
| 2521 | Construction of pAMCK12 (pXG10-bap-pRSF1010-ori-tetA) | GCGATAGACTGTATGTAAACATTTGATATCGAGCTCGCTTGG |
| 2522 | Construction of pAMCK12 (pXG10-bap-pRSF1010-ori-tetA) | CCAAGCGAGCTCGATATCAAATGTTTACATACAGTCTATCGC |
| 2523 | Construction of pAMCK12 (pXG10-bap-pRSF1010-ori-tetA) | AAAGCTTATCGATGATAAGCTGTCAAACATGAGAAAGAAGGCCATCCTGACGGATGGCC |
| 1560 | Construction of pAMCK12 (pXG10-bap-pRSF1010-ori-tetA) | TTCTCATGTTTGACAGCTTATCATCG |
| 1454 | Construction of pAMCK12 (pXG10-bap-pRSF1010-ori-tetA) | GGCTGCTGGAGATGGCGGAC |
| 2572 | Construction of pAMCK15 control (removal of sRNA44 from pAMCK14) | ATACTGAGCACTCTAGAGGCATCAAATAAAACG |
| 2573 | Construction of pAMCK15 control (removal of sRNA44 from pAMCK14) | GCCTCTAGAGTGCTCAGTATCTTGTTATCCGC |
| 2574 | Repair of mutations in sfgfp gene that originated from pXG10 bap in pAMCK12 | CCAACACTTGTCACTACTCTCACTTATGGTG |
| 2575 | Repair of mutations in sfgfp gene that originated from pXG10 bap in pAMCK12 | TTGGTGAGAATATTTGTCCTACTCAGGAGAGCGTTCACCG |

| | | |
|---|---|---|
| 2576 | Repair of mutations in sfgfp gene that originated from pXG10 bap in pAMCK12 | GACAAATATTCTCACCAATAAAAAACGCCCGGC |
| 2577 | Repair of mutations in sfgfp gene that originated from pXG10 bap in pAMCK12 | AGTAGTGACAAGTGTTGGCCATGGAACAGGTAG |
| 2578 | Repair of mutations in sfgfp gene that originated from pXG10 bap in pAMCK12 | GAGTAGGACAAATCCGCCGCCCTAGACCTAGGGTACGGG |
| 2577 | Repair of mutations in sfgfp gene that originated from pXG10 bap in pAMCK12 | AGTAGTGACAAGTGTTGGCCATGGAACAGGTAG |
